# A cationic lipid site at the outward transmembrane face of a pentameric ligand-gated ion channel

**DOI:** 10.1101/2021.03.24.436810

**Authors:** Akshay Sridhar, Sarah C.R. Lummis, Diletta Pasini, Aujan Mehregan, Marijke Brams, Kumiko Kambara, Daniel Bertrand, Erik Lindahl, Rebecca J. Howard, Chris Ulens

## Abstract

Pentameric ligand-gated ion channels (pLGICs) are crucial mediators of electrochemical signal transduction from bacteria to humans. Lipids play an important role in regulating pLGIC function, yet the structural basis for specific pLGIC-lipid interactions remains poorly understood. The bacterial channel ELIC recapitulates several properties of eukaryotic pLGICs, including activation by the neurotransmitter GABA and sensitivity to lipids, offering a simplified model system for structure-function studies. In this study, functional effects of non-canonical amino acid substitution of W206 at the top of the M1-helix, combined with detergent interactions observed in recent X-ray structures, are consistent with this region being the location of a lipid binding site on the outward face of the ELIC transmembrane domain. Coarse-grained and atomistic molecular dynamics simulations revealed preferential binding of lipids containing a positive charge, particularly involving interactions with residue W206 consistent with cation-π binding. Polar contacts from the principal subunit, particularly M3 residue Q264, further supported lipid binding via headgroup ester linkages. Aromatic residues were identified at analogous sites in a handful of eukaryotic family members, including the human GABA_A_ receptor subunit ɛ, suggesting conservation of relevant interactions in other evolutionary branches. Further mutagenesis experiments indicated that mutations at this site in ɛ-containing GABA_A_ receptors can change the apparent affinity of the agonist response to GABA, consistent with a potential role of this site in channel gating. In conclusion, this work is a detailed case study in type-specific lipid interactions at an evolutionarily distinctive pLGIC site, with implications for lipid modulation and lipophilic drug design.

## Introduction

Pentameric ligand-gated ion channels comprise a superfamily of ion channels known for their characteristic roles in fast synaptic transmission in the central and peripheral nervous systems. Elucidating the relationship of these proteins’ structures to their function has been of interest for several decades. In recent years, several atomic resolution structures from this superfamily of ion channels have been solved including members of the nicotinic acetylcholine receptors (1–4), 5-HT_3_ receptors (5–7), glycine receptors (8, 9), GABA_A_ receptors (10–15), the glutamate-gated chloride channel from *C. elegans*, GluCl (16, 17), and the prokaryote channels ELIC (18) and GLIC (19, 20). Together, these structures provide an intriguing avenue for the development of new therapeutics.

The common architectural fold observed in these structures consists of a pentameric assembly of either identical (homopentamers) or non-identical (heteropentamers) subunits each containing an extracellular domain (ECD), formed by an arrangement of β-strands connected via loops, followed by a transmembrane domain (TMD) comprised of four membrane-spanning helices (M1-M4), and an intracellular domain (ICD), which is formed by the loop connecting M3 and M4. The channel pore is lined by M2, allowing for the selective flux of permeant ions in the open conformation of the channel. The structural elucidation of representative members of this ion channel superfamily in different conformations has provided invaluable insight into the conformational changes of these channels during gating, but the role of the pore-forming transmembrane domain’s interactions with the membrane environment has been little explored (21–23).

In the past, lipids were thought to mainly serve as a structural scaffold for protein stability with occasional molecules bound, but as our knowledge of integral membrane proteins increases rapidly, so does our appreciation for the allosteric effects imparted by their immediate lipidic environments. Integral membrane proteins, including pLGICs, are organized into a specific three-dimensional structure that is governed in part by the energies associated with harmonizing the orientations of hydrophobic and hydrophilic residues with that of the host lipid matrix. It is well understood that to achieve the most energetically stable state, the length of the membrane-spanning domain containing hydrophobic residues should complement the thickness of the hydrophobic part of the lipid bilayer. This dynamic is believed to play a role in determining the organization of integral membrane proteins within the membrane (24–26).

Further insight into the functional role of lipids comes from studies on G protein coupled receptors, where negatively charged lipids enhance the receptor activation in the absence of a bilayer and in a dose-dependent manner (27); or where linker residues mediate interactions with membrane-bound cholesterols (28). More generally, as a concept that provides basis for lipids that are not just a silent surrounding for membrane proteins, evidence shows that lipids represent key factors that can influence the folding and function of different classes of membrane proteins, including ion channels, aquaporins, GPCRs, transporters and pumps (29–32).

Additionally, while it is understood that the interfacial region between membrane and water influences interactions with proteins, the effects of this interface on protein folding and binding remained poorly understood until 3D structures of bacterial porins and beta-barrel membrane proteins revealed a conserved architecture composed of a chain of aliphatic residues stretching the span of the lipid bilayer with a belt of aromatic residues, namely tryptophan (Trp) and tyrosine (Tyr), flanking the region on either end and interacting favorably with the lipid headgroups (24–26). A later biophysical approach reinforced this notion by studying the interaction of model peptides varying in hydrophobic length, but flanked by Trp or lysine (Lys) residues at both ends, and their influence on lipid bilayers. They found that upon decreasing the peptide’s hydrophobic length, the lipids adopted more nonlamellar phases; alleviating hydrophobic mismatch, and illustrating the preferred membrane-anchoring effect of Trp and Lys residues (33).

In this study, we focused our attention to such a Trp residue flanking the top of the M1-helix in the transmembrane domain of ELIC, a model prokaryote pLGIC. Using a combination of complementary techniques, we investigate the functional contribution of this Trp residue to channel gating. Employing unnatural amino acid mutagenesis and electrophysiological recordings, we demonstrate that the Trp residue is involved in a cation-π interaction. Together with the known ability of the Trp residue to interact with the polar head group of a detergent molecule, we further investigate whether the Trp residue interacts with cationic or zwitterionic lipids using molecular dynamics simulations. Finally, we extrapolate these findings to GABA_A_ receptors containing a conserved aromatic residue at this position. Together, our work sheds new light on lipid interactions in certain pLGICs.

## Results

### Residue W206 at the outward face of the ELIC transmembrane domain is involved in a cation-π interaction

To investigate the functional importance of tryptophan residues in the gating of ELIC, we focused on residue W206, which is near the boundary between the extracellular ligand binding domain and the pore-forming transmembrane domain. This site at the top of the M1-helix forms part of the outward face of the ELIC transmembrane domain, which contacts the lipid bilayer. In ELIC, the W206 side chain points outward and into the lipid bilayer, potentially forming interactions with lipids. Recently, we determined an ELIC structure bound to a nanobody at a resolution of 2.5 Å, thus revealing new details of interacting ions, lipids and detergent molecules. In this structure, W206 interacts with an undecylmaltoside molecule, with the W206 side chain pointing toward the polar head group and the lipophilic tail pointing downward along the M1-helix (Figure 1A). Notably, this site is homologous to the ivermectin binding site in the glutamate-gated chloride channel from *C. elegans* (Figure 1B, pdb accession code 3rif) (16) and a POPC lipid binding site in the α1β3γ2 GABA_A_R structure (Figure 1C, pdb accession code 6i53) (13). These observations raised the question as to whether the W206 in ELIC could also interact with lipid molecules in the context of the native lipid bilayer. Strikingly, an aromatic residue at this position is conserved in selected eukaryote receptors, including a putative 5-HT_3_R in *C. latens*, the ɛ GABA_A_R subunit in *Homo sapiens*, different insect GABA_A_R subunits (*Papillio machaon*, *Heliothes viricens*, *Aedes aegypti*), nAChR subunits from coral (*Acropora digitifera*, *Acropora millepora* and *Stylophora pistillata*), the histamine-gated chloride channel from *Drosophila* and various nAChR subunits from *Drosophila* and *Apis mellifera*. This conservation of an aromatic residue suggests a possible structural or functional role in these receptors.

**Figure 1.**
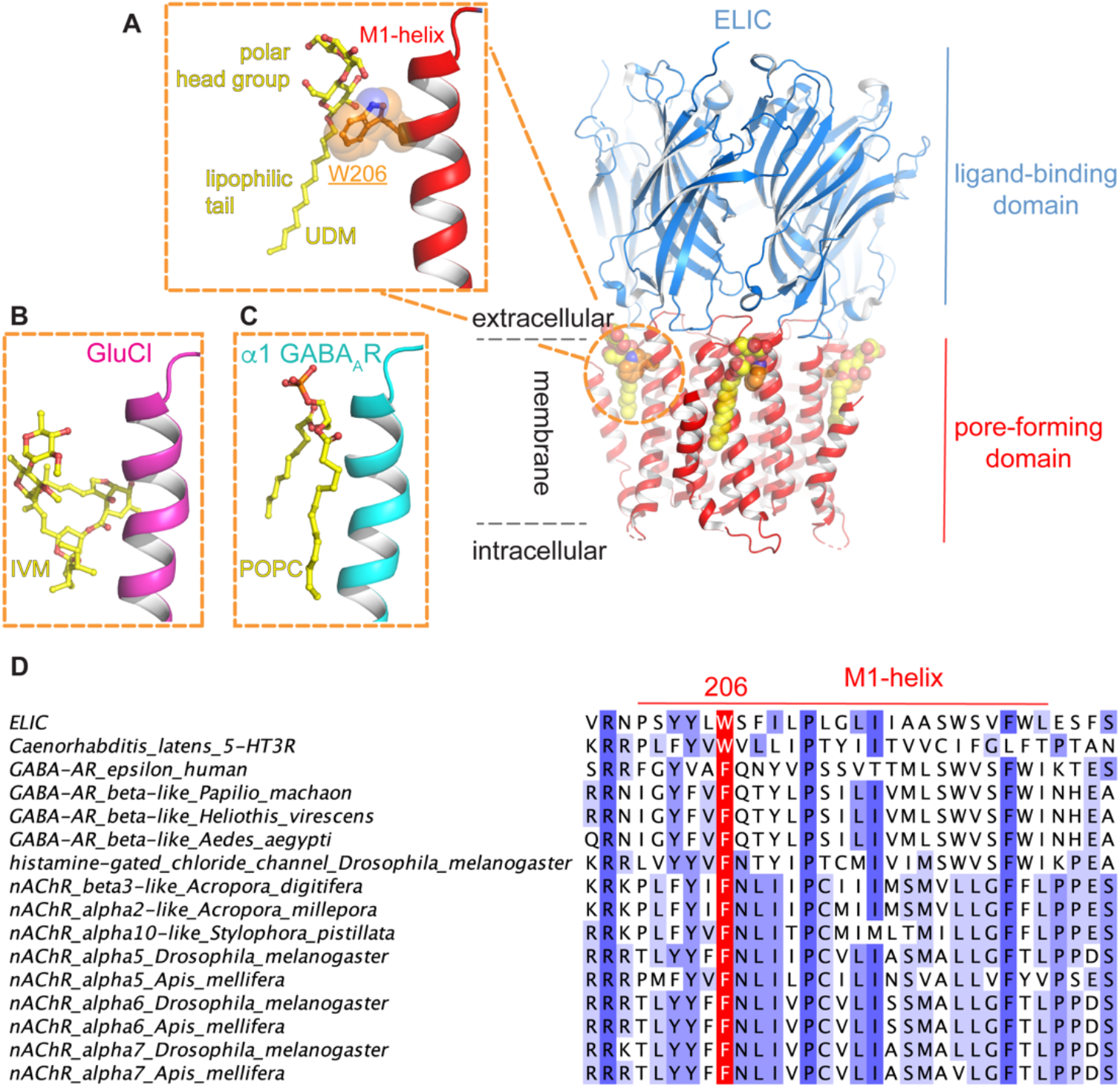
**A**. Location of a putative lipid binding site at the outward transmembrane face of the pentameric ligand-gated ion channel ELIC. ELIC is shown in cartoon presentation with the extracellular ligand-binding domain shown in blue and the pore-forming transmembrane domain in red. The putative lipid binding is located near W206, shown as orange sticks, in the M1-helix. In a recently published 2.5 Å ELIC structure (pdb accession code 6hjx) (22) this site is occupied by a detergent molecule (undecylmaltoside), shown in yellow spheres. The inset shows a detail of the interaction, with the polar head group forming an interaction with W206. **B**. This binding site is homologous to the ivermectin (IVM, shown in yellow sticks) binding site in the glutamate-gated chloride channel from *C. elegans*, GluCl (pdb accession code 3rif) (16) and (**C**) a POPC (yellow sticks) binding site in the α1β3γ2 GABA_A_R structure (pdb accession code 6i53) (13). **D**. Sequence alignment showing conservation of aromatic residues (colored in red) at the position homologous to W206 in ELIC. Residues are colored in shades of blue by using an identity threshold of 20%.

To examine the contribution of W206 in this interaction, we investigated wild type (WT) and mutant ELIC, transfected them into HEK293 cells, and probed GABA-elicited responses in a FlexStation using membrane potential sensitive dye (Figure 2A–B). Concentration-response curves for wild type receptors revealed a GABA EC_50_-value of 1.1 mM (pEC50 = 2.96 ± 0.1) and a Hill coefficient of 2.2 ± 1.4, consistent with previously published data. Alanine substitution of W206 resulted in nonfunctional receptors (no response with up to 100 mM GABA) but we observed robust responses in receptors with W206Y (EC_50_ 5.3 mM, pEC_50_ = 2.272 ± 0.09, n=4) and W206F mutations (EC_50_ 2.0 mM, pEC_50_ = 2.694 ± 0.04, n=4). pEC_50_-values of these receptors were increased compared to wild type (p=0.0008), and, for W206Y-containing receptors, maximal responses were smaller (p=0.0013). To determine if the π ring contributes to the interaction of W206, we substituted fluorinated Trp residues using non canonical mutagenesis followed by two-electrode voltage clamp of expressed receptors in oocytes (Figure 2C). All the fluorinated Trps resulted in increased EC_50_-values and plotting relative EC_50_s against cation-π binding energy demonstrated a good correlation (r2= 0.98, Figure 2D) indicating that such an interaction is important here for the function of the receptor.

**Figure 2.**
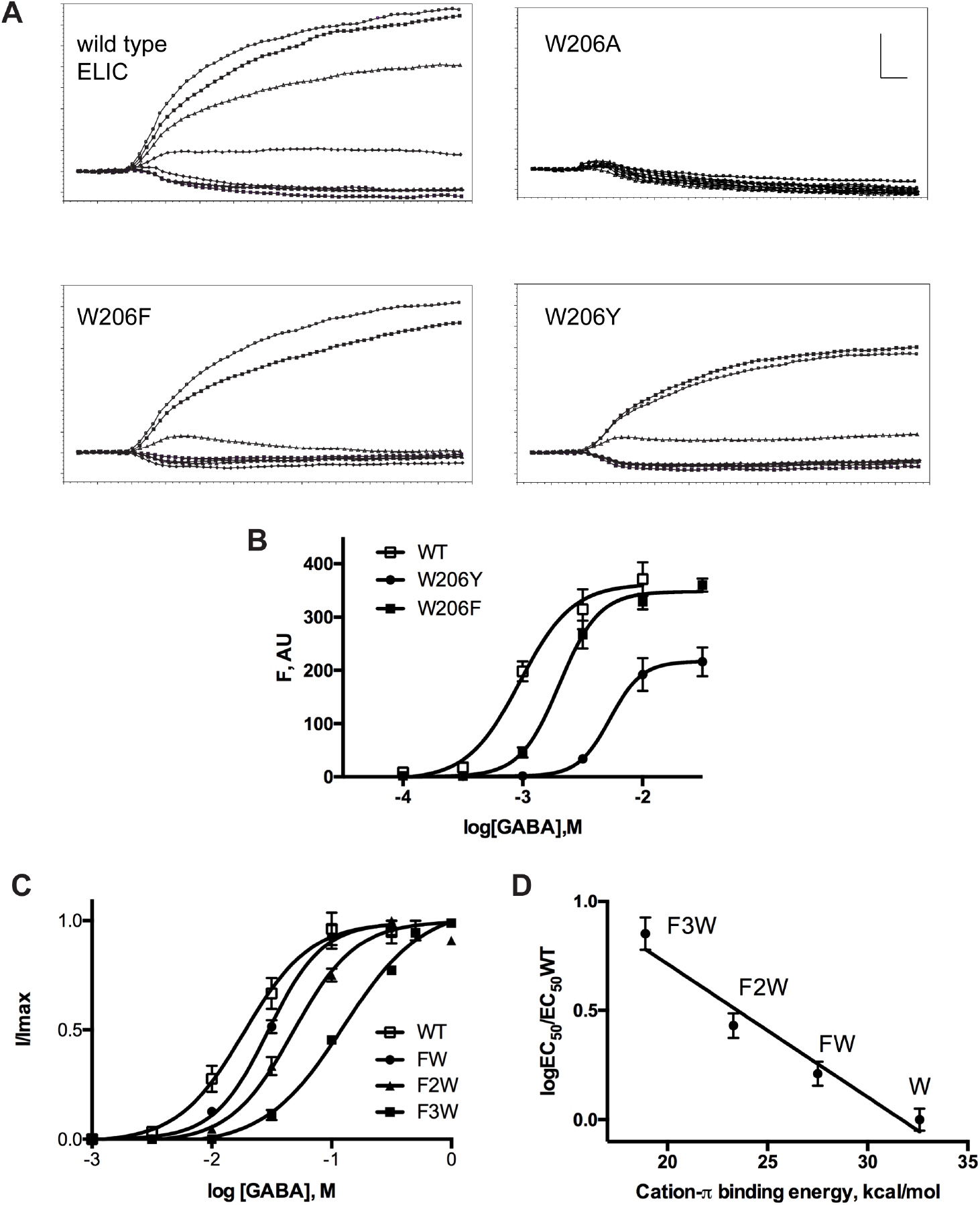
**(A**) Typical FlexStation responses to application of GABA (0, 0.1, 0.3, 1, 3, 10, and 30 mM) at 20 s to HEK293 cells transfected with wild type and mutant ELIC. Scale bar = 100 F, AU (arbitrary units) and 10 s. **(B)** Concentration-response curves from FlexStation data (mean ± SEM, n=4). **(C)** Concentration-response curves for ELIC assayed in oocytes showing the effects of incorporation of the noncanonical amino acids FW, F2W and F3W at position 206 (mean ± SEM, n=4). **(D)** Fluorination plot of W206 in ELIC. EC_50_-values for ELIC activation by the agonist GABA in wild type (WT) and 1-F, 2-F, and 3-F substituted W206 are indicated as FW, F2W and F3W, respectively. The plot of the EC_50_-values relative to the cation-π binding energy reveals a linear correlation (r2= 0.98), which is indicative of a strong cation-π interaction with W206 in ELIC.

### An aromatic lipid-binding site at the outward complementary face of the ELIC transmembrane domain

To identify the specific lipid interactions contributing to the observed role of the W206 residue, we first performed triplicate 10-μs molecular dynamics simulations using the coarse-grained MARTINI model (34). To differentiate on the basis of charge, a test membrane containing 20% anionic palmitoyloleoylphosphatidylglycerol (POPG) and 80% zwitterionic palmitoyloleoylphosphatidylcholine (POPC) lipids in each leaflet was simulated around the restrained protein. Consistent with previous reports (23), preferential interactions of POPG were observed in the inner leaflet, involving a cluster of basic residues on the inward-facing M3-M4 loop (Figure 3A). The more prevalent POPC was involved in a larger number of interactions, partly at the inward-facing M3-M4 site, but also an outward-facing site at the junction of the extracellular and transmembrane domains, involving the pre-M1 motif and upper M1 helix -including residue W206- as well as proximal sites in upper M3 and M4 (Figure 3B).

**Figure 3.**
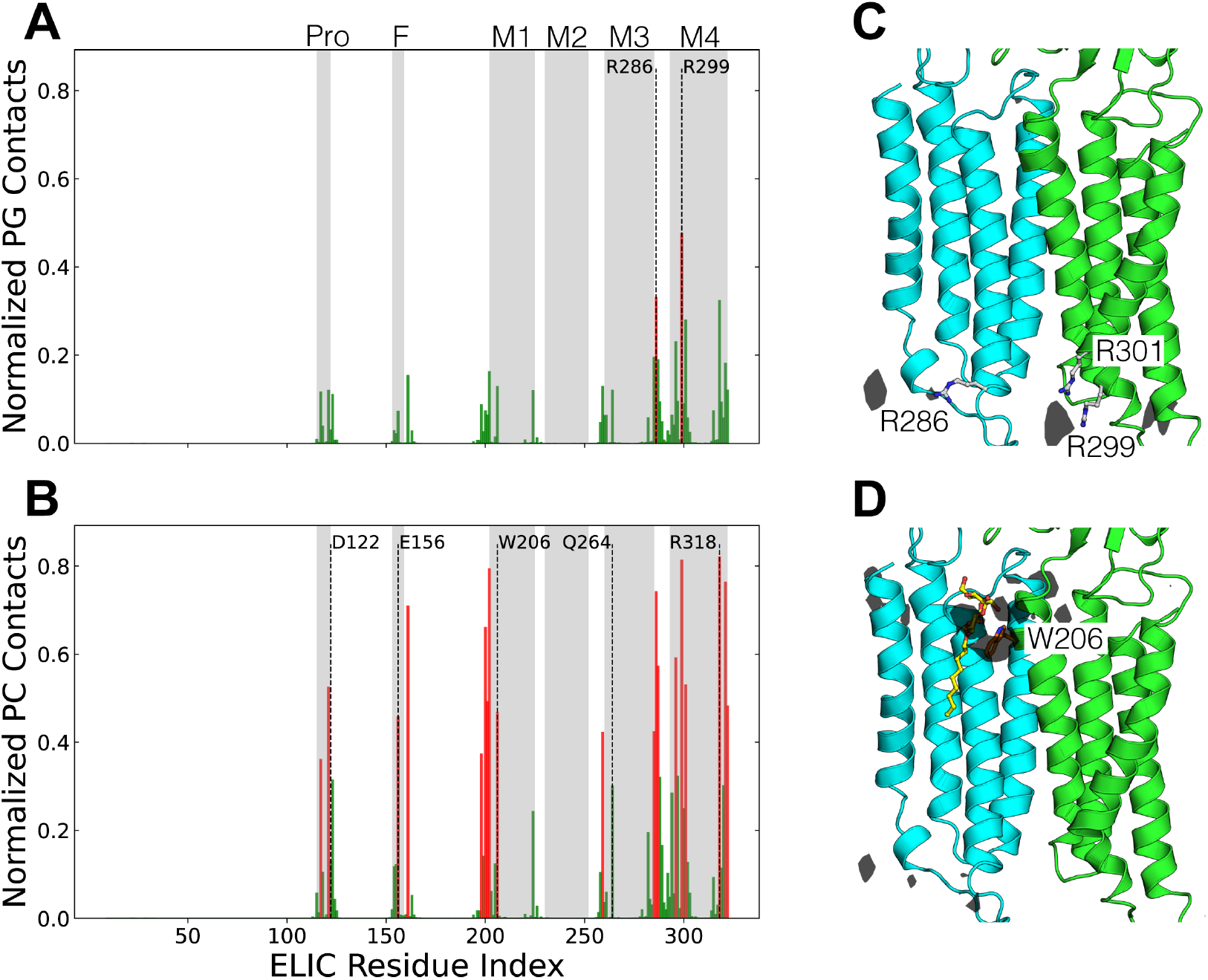
**(A,B)** Normalized number of contacts between ELIC residues and headgroups of the POPG/POPC lipids from coarse-grained simulations. A contact was assumed if a residue’s bead was within 5.5 Å of the lipid headgroup’s bead and the contacts were subsequently averaged across the five subunits. Residues with a contact frequency of >33% are colored red. Residue indices corresponding to structural elements are highlighted in grey and important residues within them are highlighted. **(C,D)** Densities of the different PG/PC lipids calculated from CG simulations illustrated at one of the subunit interfaces at an isosurface value of 1.9 molecules/nm^3^ and 5.0 molecules/nm^3^, respectively. For POPG, the R286, R299 and R301 residues mediating interactions with the anionic lipid are illustrated. For POPC, the M1-helix W206 residue adjacent to the preferential residence site is illustrated. Additionally, the detergent molecule identified to bind at this site in the PDB structure 6hjx is overlaid on the subunit interface and illustrated in yellow.

We further investigated the structural basis for specific interactions in this putative outer-leaflet lipid site using unrestrained all-atom molecular dynamics simulations. As a starting model, ELIC was backmapped to atomistic resolution along with a representative POPC molecule at each of the five putative W206 sites. In triplicate simulations launched from this configuration, targeted POPC molecules were relatively stable, with the middle 50% between 6 and 8 Å root mean-squared deviation (RMSD) from the starting pose (Figure 4A). Moreover, POPC headgroups distributed with a median distance <6 Å from the W206 sidechain, characteristic of a cation-π interaction (Figure 4B) (35). In contrast, substituting alanine for the central tryptophan (W206A) allowed the lipid to deviate more widely, with the middle 50% ranging 8-13 Å in RMSD (Figure 4A) and occupying a spread of positions centered >8 Å from the mutated residue (Figure 4B). Thus, W206 appeared to be important in retaining POPC at the outer-leaflet site. To test the charge dependence of this apparent interaction, we also ran atomistic simulations with the cationic lipid 1,2-dipalmitoyl-3-trimethylammonium-propane (DOTAP) or anionic POPG in place of zwitterionic POPC at each interface. Whereas behavior of DOTAP was markedly similar to POPC, anionic POPG deviated further from its starting position (Figure 4A) and occupied a broad range of distances centered >9 Å from W206 (Figure 4B), consistent with cation dependence.

**Figure 4.**
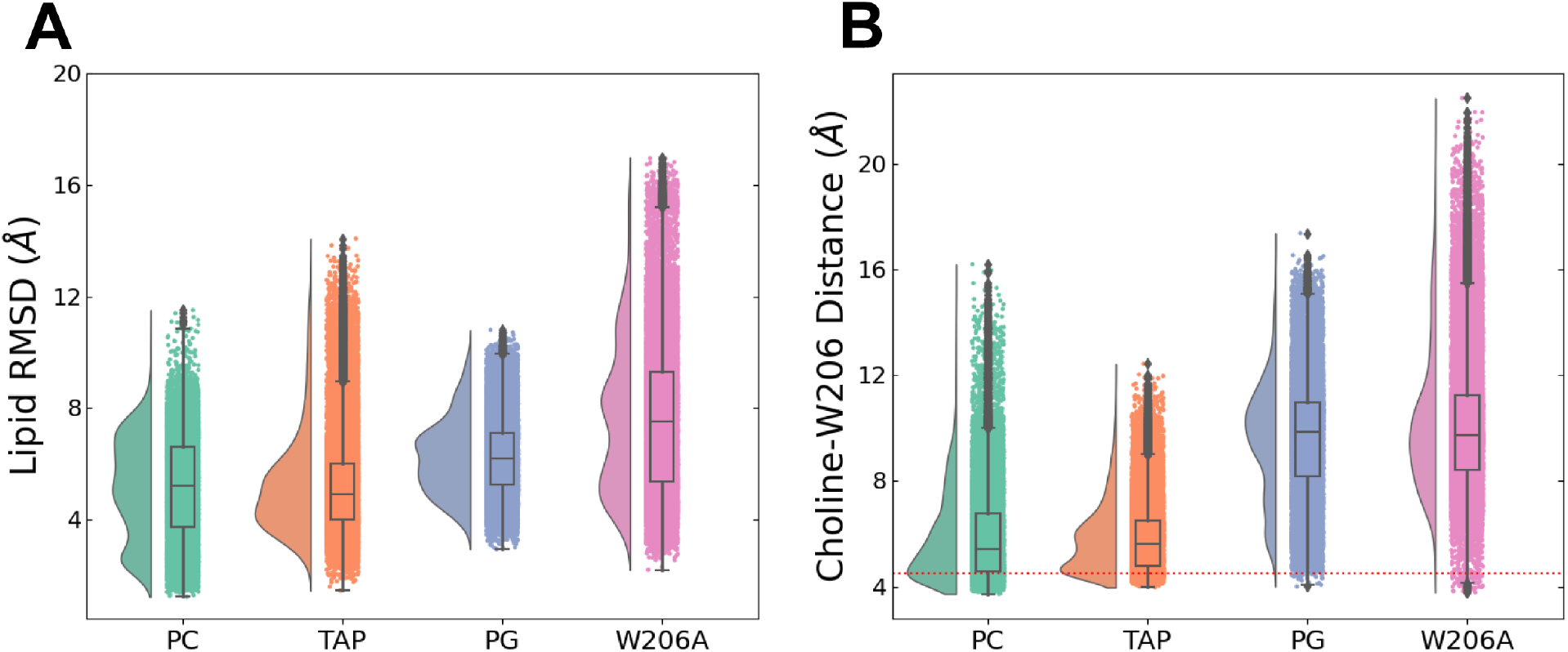
**(A)** Stability of various lipid types at the inter-subunit binding site illustrated by the RMSD probability distributions and box plots. For this calculation, the Cα atoms of the M2 helix were first aligned and the RMSD was calculated for the non-hydrogen lipid head-group atoms. The RMSD values were then averaged across the five subunit interfaces over the final 150 ns of three independent 300 ns MD trajectories. **(B)** The role of cation-π interactions in stabilizing the lipid at the subunit interface illustrated by the distribution of distance between the choline lipid headgroup and the center of the W206 aromatic sidechain. For each lipid type, the distances were averaged across the five subunit interfaces over the final 150 ns of 3 independent 300 ns MD trajectories. For the PG lipid without the choline group, the distances were instead calculated to the phosphorus atom and the ideal cation-π binding distance of 4.5 Å is illustrated as a dotted red line.

### Polar lipid contacts on the principal subunit

To elucidate structural determinants of lipid binding in the upper-leaflet site, we also probed contacts on the principal neighboring subunit (Figure 5). Atomistic simulations revealed prolonged interactions of M3-Q264 and M4-R318 with lipid-headgroup ester linkages (Figure 5A–C). Q264 was previously seen to coordinate detergent molecules bound at W206 (22); a role for R318 was less definitive, as this residue is not well resolved in many reported structures, and was modeled *ab initio* for MD simulations. Moreover, removal of the M4 arginine sidechain (R318A) was previously shown to enhance rather than diminish channel activity (36); indeed, ELIC has been shown to retain channel activity even upon deletion of M4 (22), suggesting the M4-lipid interaction is not critical to function. To test the relevance of these apparent contacts, we ran additional simulations in the presence of DOTAP with removal of the M3-glutamine sidechain (Q264A), the entire M4 helix (ΔM4), or both features (Figure 6). Either Q264A or ΔM4 allowed the lipid to deviate further from its starting pose than in the wild-type system, and their combined effect was greater than either individual modification (Figure 6A). Moreover, analysis of choline-W206 distances indicated considerable dissociation of this cation-π interaction in the presence of Q264A, with or without M4 (Figure 6B). Thus, polar contacts in M3 and possibly M4 appeared to directly support cation-π lipid interactions with the complementary M1-helix.

**Figure 5.**
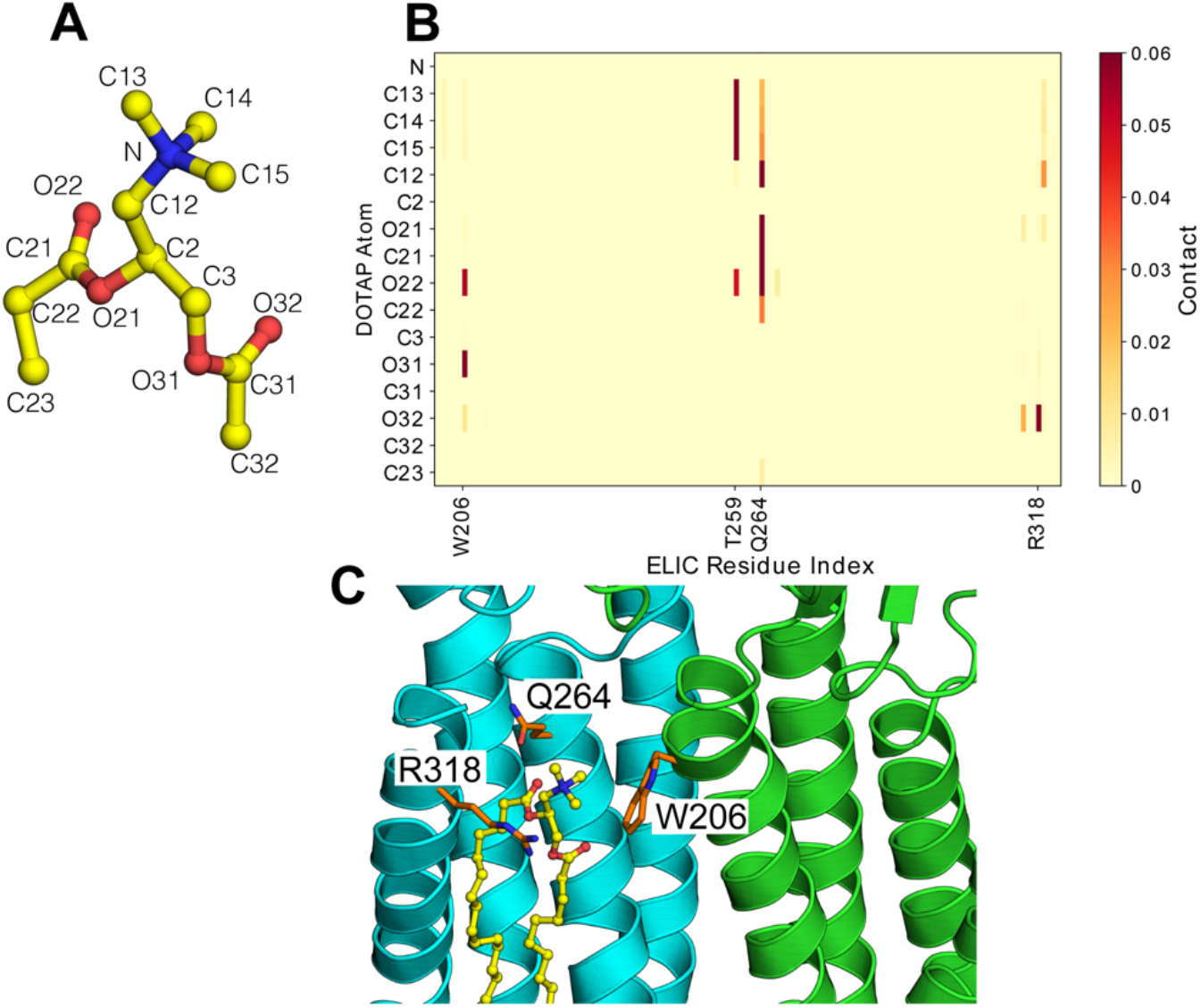
**(A,B)** Secondary contacts stabilizing the lipid at the inter-subunit binding site illustrated as a contact map with the headgroup of the cationic DOTAP lipid. A contact was assumed if a non-hydrogen atom of the residue was within 3.2 Å of the lipid atom. The contacts are averaged across the five subunit interfaces over the final 150 ns of three independent 300 ns MD trajectories. **(C)** The W206, Q264 and R318 side chains are illustrated in orange relative to the position of the lipid (in yellow) at the subunit interface.

**Figure 6.**
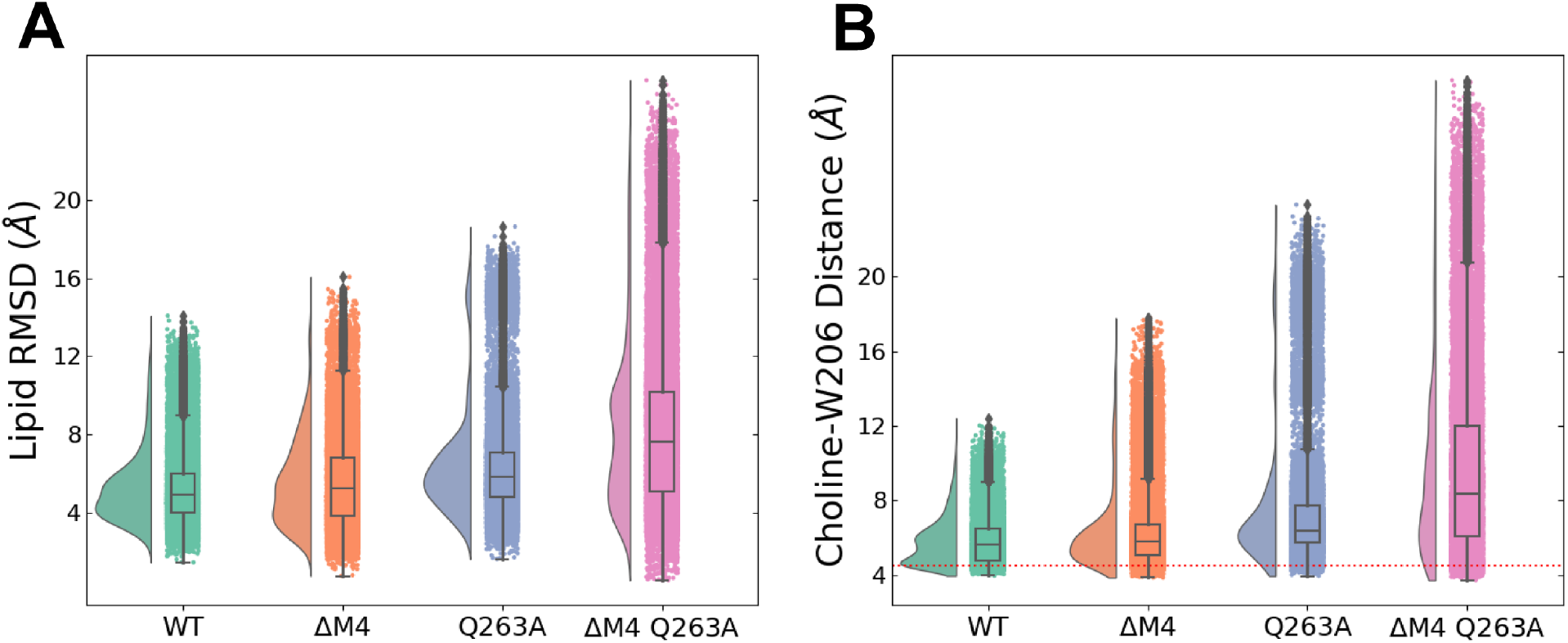
**(A)** The role of the M2 and M4 polar contacts in stabilizing the lipid at the subunit interface analyzed by lipid RMSD probability distributions and box plots. For this calculation, the Cα atoms of the M2-helix were first aligned and the RMSD was calculated for the non-hydrogen DOTAP head-group atoms. The RMSD values were then averaged across the five subunit interfaces over the final 150 ns of three independent 300 ns MD trajectories. **(B)** The role of the M2- and M4-helix polar contacts in stabilizing the lipid at the subunit interface analyzed by the distribution of the DOTAP Choline-W206 sidechain distance. The distances were averaged across the five subunit interfaces over the final 150 ns of 3 independent 300 ns MD trajectories. The ideal cation-π binding distance of 4.5 Å is illustrated as a dotted red line.

### Site-directed mutagenesis of M1 aromatic residue in α1β2ɛ GABA_A_R expressed in *Xenopus* oocytes

To verify the importance of an aromatic residue at the top of the M1-helix in eukaryote receptors, we mutated the homologous residue in the ɛ-subunit GABA_A_R subunit, F260 (numbering according to mature protein). Using two electrode voltage clamp recordings from *Xenopus* oocytes expressing wild type and mutant receptors, comprising ɛ plus α1 and β1 subunits, we then determined EC_50_-values in response to the agonist GABA. Substitution of a nonaromatic glutamic acid residue (ɛ F260E) decreased apparent GABA affinity 3-fold (Figure 7): GABA EC_50_-values were 2.27 ± 0.27 (n=6, wild type) and 6.52 ± 0.54 (n=12, ɛ F260E). A t-test indicates a *P*-value of < 2.5 ×10^−6^. This disruptive effect was consistent with a direct or indirect role of F260 in gating of ɛ-containing GABA_A_Rs, possibly involving lipid modulation, although insignificant effects of other substitutions (F260I, F260R) may indicate compensatory or complex interactions at this site.

**Figure 7.**
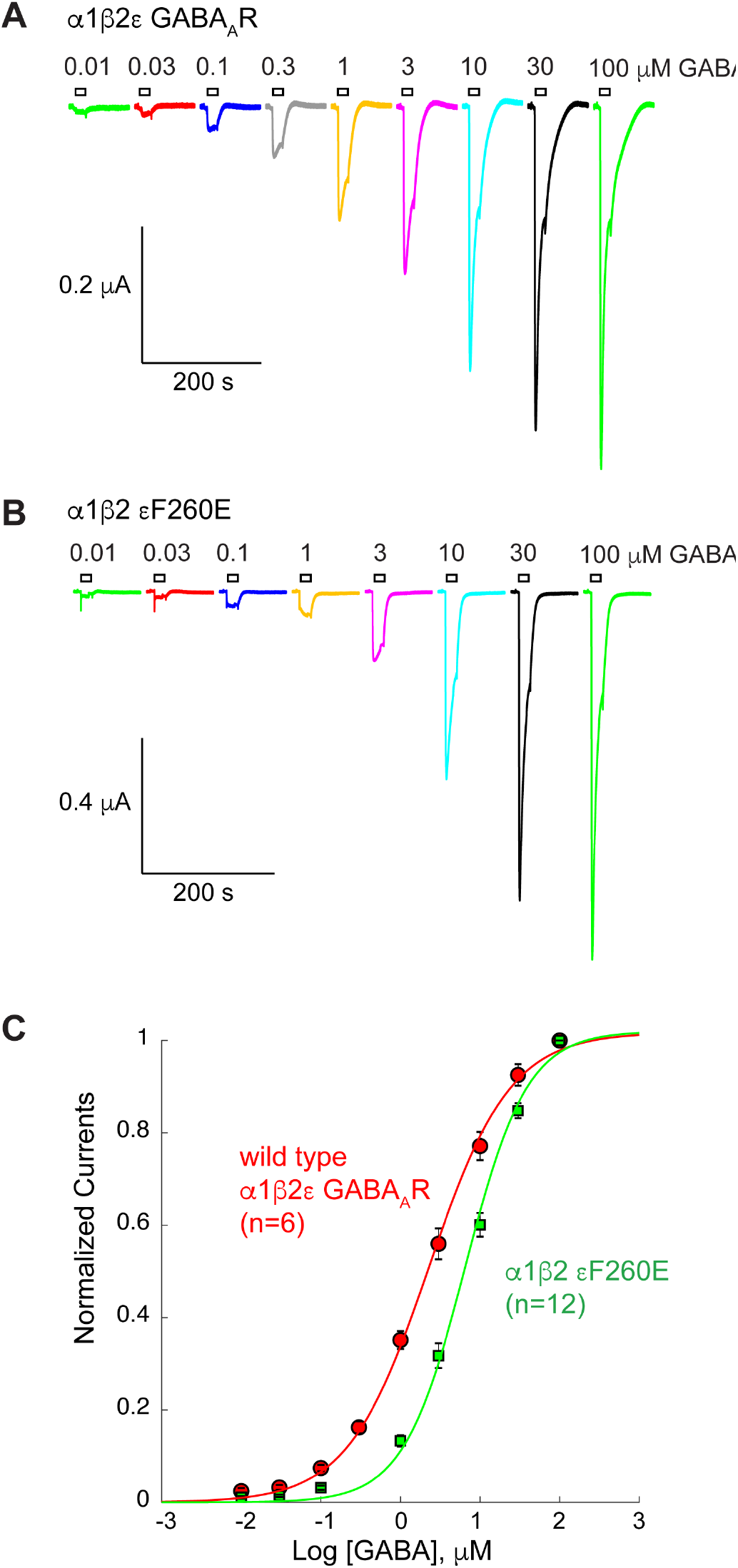
**(A,B)** Representative current traces evoked from *Xenopus* oocytes expressing the wild type human α1β2ɛGABA_A_R (**A**) and the α1β2 ɛF260E mutant (**B**). Agonist-evoked currents were obtained by application of the GABA concentrations as indicated. (**C**) Concentration-response relationships for the experiments indicated in A and B. GABA EC_50_-values were 2.27 ± 0.27 (n=6, wild type) and 6.52 ± 0.54 (n=12, ɛ F260E). A t-test indicates a *P*-value of < 2.5 ×10^−6^.

## Discussion

An increasing number of structural and biophysical studies have revealed lipid interactions in different classes of membrane proteins, including ion channels and GPCRs, which are important drug targets (reviewed in 37, 38). These lipids can either fulfill a structural role, through binding at a defined location in the transmembrane domain, or can additionally influence protein function through allosteric effects. Importantly, molecular dynamic simulations have played a crucial role in identifying certain lipid interactions (37), through complementation of experimental structural studies, such as X-ray crystallography or cryo-EM, which often reveal electron densities that do not permit an unambiguous assignment of the lipid identity.

One of the best-described examples comes from the class of inwardly rectifying K^+^ (Kir) channels, which are activated by the anionic phospolipid PIP2 (phosphatidylinositol 4,5-bisphosphate) (38). The PIP2 interaction site has been observed in several Kir structures, including Kir2 (39), and involves highly conserved basic residues (arginine and lysine) from the TM helices and cytoplasmic domain that interact with the negatively charged PIP2. Combined electrophysiological studies and MD simulations have revealed the dynamic nature of the PIP2 interaction (40), which involves a stabilization of the interaction between the TM domain and the cytoplasmic domain, thereby opening the channel gate.

In the related class of voltage-gated (Kv) K^+^ channels, a phosphatidylglycerol (PG) lipid has been observed in the crystal structure of a Kv1.2/Kv2.1 chimeric channel. Here, the lipid wedges between the S1-S4 voltage-sensor domain (VSD) and the S5-S6 pore domain, thereby coupling voltage-sensor motions to pore opening (41). A combination of MD simulations (42) and functional studies (43) have indeed confirmed the phospholipid interactions with arginines in the voltage sensor and indicated that phospholipids facilitate voltage-driven conformational transitions, thereby enabling lipid-dependent gating of Kv channels (44).

In pLGICs, a prominent role of the M4-helix has emerged in lipid sensing as it directly contacts the lipid bilayer (21, 22). In a recent study on ELIC, we demonstrated that a phosphatidylethanolamine (PE) lipid binds at the lower half of the M1- and M4- helices and at a nearby site for neurosteroids (45–47), cholesterol (11) and general anesthetics (48). This site is shaped by a characteristic proline-kink halfway the M4-helix, which is conserved in eukaryote GABA_A_ and glycine receptors. Using a combination of complementary methods, we demonstrated that M4 is intrinsically flexible and that M4 deletions or mutations of the lipid binding site accelerate desensitization, a phenomenon that can be mimicked by reconstitution of ELIC into membranes of different lipid composition (22). These data indicate that M4 may act as a lipid sensor and that lipid interactions shape the agonist response. Further evidence into the role of the M4-helix comes from studies on the *Torpedo* nAChR, which revealed that in the absence of anionic lipids and cholesterol, the receptor binds agonist, but does not undergo agonist-induced conformational transitions, a phenomenon that has been called receptor “uncoupling” (21).

In addition to the lipid binding site at the lower half of the M4-helix, several other lipid binding sites have been resolved in pLGICs. A detailed review of each of those individual lipid sites would be beyond the scope of this discussion, but as an example in GLIC, two phosphatidylcholine (PC) binding sites are located in a groove between M4 and both M1 and M3 (19), one in the upper half and the other in the lower half of the TM domain. The lipid bound in the upper half of the TM domain is displaced in the GLIC structure bound to the general anaesthetic propofol (48).

As an example in eukaryote receptor structures, a cryo-EM structure of the full length α1β3γ2 GABA_A_R in lipid nanodiscs revealed PIP2 molecules bound to the intracellular side of the TM domain, forming polar interactions between the PIP2 phosphate headgroup and basic (arginine and lysine) sidechains of the α1 M3 and M4 helices (13). In a recent cryo-EM structure of the α4β2 nAChR, two cholesterol molecules per receptor subunit were bound at the receptor periphery along the intracellular half of the transmembrane domain and flanking the subunit interface (2).

Within the context of the present study, which focuses on a lipid binding site at the top of the M1-helix (W206 in ELIC) it is worth discussing several other studies that have revealed lipids or lipophilic compounds bound at or near to this site in different pLGICs. For example, in the glutamate-gated chloride channel from *C. elegans* (pdb code 3rif) (16), the allosteric agonist ivermectin is bound to the same site, forming interactions with L218 (equivalent to W206) and V278 (equivalent to Q264). In the α1β3γ2 GABA_A_R structure (pdb code 6i53) (13), POPC is in contact with α1-I228 (equivalent to W206) and β3-M283 (equivalent to Q264), although the lipid is somewhat excluded from the cleft. In α1β2γ2 GABA_A_R structures (pdb codes 6×3×, 6×3t) (15), weak putative lipid densities are intercalated at α1/β2 and α1/γ2 interfaces, potentially proximal to β2-L223/ γ2-I238 (equivalent to W206) and/or α1-W288 (equivalent to Q264), although lack of resolution precludes identifying direct contacts. In 6×3t, propofol is resolved at β2/α1 interfaces, deeper than ELIC lipids but still in contact with the backbone of α1-I228 (equivalent to W206), and enabling visualization of additional putative lipid densities at the interface periphery. Taken together, these studies demonstrate that the W206 site at the top of the M1-helix in ELIC is structurally equivalent in different pLGICs and can serve as a binding site for lipids or lipophilic drugs.

In summary, our study investigates the outward transmembrane face in ELIC as a lipid binding site. Using a combination of complementary methods, we investigated the role of W206 in lipid interactions. Using non-canonical mutagenesis we reveal that W206 is involved in a cation-π interaction. Together with previous structural data showing that the W206 engages in interactions with the polar headgroup of a detergent molecule, we investigated interactions of W206 with cationic and zwitterionic lipids using molecular dynamics simulations. Using site-directed mutagenesis we show that the conserved aromatic residue in the GABA_A_R ɛ-subunit affects gating. Together, these results expand our knowledge of lipid interaction sites in pLGICs.

### Experimental procedures

#### FlexStation™ methods

These methods were similar to those previously described (49). Briefly, ELIC cDNA was transfected into HEK293 cells and these were then grown for 2-3 days in a 96-well plate. Then blue fluorescent membrane potential dye (Molecular Devices Ltd., Wokingham, UK) diluted in Flex buffer (10 mM HEPES, 115 mM NaCl, 1 mM KCl, 1 mM CaCl_2_, 1 mM MgCl_2_, 10 mM glucose, pH 7.4) was added to each well. After incubation at 37°C for 30 min, plates were placed in a FlexStation^™^ (Molecular Devices Ltd.) and fluorescence measured every 2 s for 120 s. Buffer or GABA (0.03-30 mM) was added to each well after 20 s. Concentration-response data were fitted to the four-parameter logistic equation, F = F_min_ + (F_max_ − F_min_)/(1+10^(log(EC50-[A])*nH^), where [A] is the concentration of agonist, nH is the Hill coefficient, and F_max_ and F_min_ are the maximal and minimal fluorescence levels for each dataset, using Prism software (GraphPad, San Diego, CA). Wild type and mutant responses were compared using an ANOVA test followed by Dunnetts multiple comparison test.

### Non-canonical Amino Acid Incorporation

Site-directed mutagenesis was performed using the QuikChange strategy (Stratagene) using ELIC in pGEM-HE. Mutations were confirmed by sequencing. For non-canonical amino acid mutants the site of interest was mutated to the TAG stop codon. Plasmids were linearized and receptor mRNA prepared by in vitro runoff transcription using the Ambion T7 mMessage mMachine kit. Non canonical amino acids ligated to tRNA were prepared as previously described (50). Stage V-VI oocytes of *Xenopus laevis* were harvested and injected with mRNAs as described previously. For wild type experiments and conventional mutants, each cell received a single injection of 10-25 ng of receptor mRNA approximately 24 h before recording. For nonsense suppression experiments, each cell was injected with 50-100 ng each of receptor mRNA and appropriate tRNA approximately 48 h before recording. Injection volumes for each injection session were 50-100 nL per cell.

### ELIC electrophysiology

Two-electrode voltage clamping of Xenopus oocytes was performed using an OpusXpress system (Axon Instruments, Inc., Union City, CA). All experiments were performed at 22-25 ºC. GABA (Sigma) was diluted in ND-96 and delivered to cells via a computer-controlled perfusion system. Glass microelectrodes were backfilled with 3 M KCl and had a resistance of approximately 1 MΩ. The holding potential was −60 mV unless otherwise specified. Concentration-response curves and parameters were obtained using Prism software (GraphPad, PRISM, San Diego, CA).

### Model Building

As a starting model for ELIC simulations, protein atoms from a 2.5-Å resolution X-ray structure (PDB ID 6HJX) (22) were extracted from their associated nanobodies, lipids, and detergent molecules. Unresolved residues at the M4 C-terminus of each subunit were built as a continuous helix using PyMOL. Since cation-π interactions are due to the orbital orientations in aromatic rings not treated explicitly in simulations, care was taken to use force fields with appropriate corrections for all models (51).

### Coarse-Grained Simulations

Coarse-grained simulations were performed using the MARTINI 2.3P polarizable force field (34) with improved choline-aromatic cation-π interaction parameters (52). Each protein was embedded in a symmetric membrane containing 80% POPC and 20% POPG, or a brain-lipid mimic mixture as previously described (15), using the Martini Bilayer Maker (53) in CHARMM-GUI (54). The membrane spanned a dimension of 300 × 300 Å containing 2180 lipids and 283,597 beads including ions and polarizable water (55). After energy minimization and equilibration for 20 ns, three replicates of each system were simulated for 10 μs using GROMACS 2018 (56), with all protein beads restrained to allow convergence of lipid interactions.

The final 7.5 μs of the simulation trajectories were used for analysis using Python MDAnalysis/MDTraj scripts (57, 58). A contact was assumed if a residue’s bead was within 5.5 Å of a lipid head-group bead, and occupancy probability-density calculations were performed using a 2 Å-resolution grid.

### Atomistic Simulations

As a starting model, ELIC was backmapped to atomistic resolution using Backward (59) along with a POPC molecule from a representative coarse-grained simulation frame, occupying the high-probability volume associated with a single receptor subunit. This lipid was then replicated and symmetrized to occupy each of the five putative sites in the pentameric channel. PyMOL (http://www.pymol.org) was used to introduce mutations at the lipid binding site, and alternative lipids (DOTAP, POPG) were substituted by alignment of their head- and acetyl groups. The structure with five bound lipids was then placed in a POPC bilayer of dimension 150 × 150 Å using CHARMM-GUI (54). The Charmm36M force field (60) with WYF cation-π corrections (61) was used to describe the system. Energy minimization and equilibration were performed with restraints of 1000 kJ mol^−1^ nm^−2^ on the bound lipids.

After equilibration, three replicates of each system were simulated for 300 ns using GROMACS 2018 (56) and a timestep of 2 fs. Long-range electrostatic interactions were calculated using the particle mesh Ewald method (62) and hydrogen-bond lengths were constrained using LINCS (63). Pressure and temperature were maintained through the use of the Parrinello-Rahman barostat (1 bar) (64) and v-rescale (300 K) thermostat (65), respectively. Lipid binding stabilities were calculated as an average over all five subunit interfaces from the final 150 ns of all replicates.

### GABA_A_ Receptor Electrophysiology

The sequence of the gene encoding the GABA_A_R ɛ subunit corresponding to the accession number NM_004961 was synthesized by Blue Heron Biotech’s into the pCMV6-AC vector from Origen, which offers the advantage of allowing expression in eukaryote cells with the cytomegalovirus (CMV) promotor as well as the bacterial T7 promotor for *in vitro* synthesis of mRNA. The nucleotide sequence was optimized for expression in mammalian cells using standard procedures. Mutations were engineered with a QuikChange strategy and confirmed by sequencing. For functional expression of GABA_A_R containing the ɛ subunits the mRNA’s encoding for the human α1 (NP_000797.2), the β2 (NP_068711.1) and ɛ subunits were mixed in a ratio of 1:1:1 in nuclease free distilled water at a concentration of 0.4 μg/μL.

Following standard preparation of the oocytes (66), stage V and VI cells were manually selected under a binocular and disposed in a 96 microtiter plate previously filled with ND96-solution containing 96 mM NaCl, 2 mM KCl, 1.8 mM CaCl_2_, 2 mM MgCl_2_ and 5 mM HEPES, pH 7.4, supplemented with 50 mg/L gentamicin sulfate. Injection of 2 ng of mRNA per oocyte was done using the automated injection system Roboinject (Multi Channel Systems). Oocytes were incubated at 18°C for two to five days prior to conducting the electrophysiological recordings using the two electrode automated voltage clamp system (HiClamp apparatus, Multi Channel Systems). A standard OR2 solution containing 82.5 mM NaCl, 2.5 mM KCl, 1.8 mM CaCl_2_, 1 mM MgCl_2_ and 5 mM HEPES buffered at pH 7.4 was used as control and cells were maintained a 20°C using the cooling system of the HiClamp. Currents were evoked by brief exposure to GABA as indicated on the figures. Data acquired with the HiClamp were analyzed using the manufacturer’s software (Multi Channel Systems). Concentration-activation curves were fitted with the empirical Hill equation. Data are presented as the mean ± standard error of the mean (SEM). Statistical comparison between wild type and mutants was done with an unpaired, two-tailed t-test with Welch’s correction for unequal sample size and variance.

## Data availability

Sample frames from coarse-grained and atomistic MD trajectories are available on zenodo.org with doi 10.5281/zenodo.4618338.

## Acknowledgements

AS was supported by Marie Skłodowska-Curie grant 898762, and EL/RJH by grants from the Swedish Research Council (2017-04641, 2019-02433) and Swedish e-Science Research Center. Computational resources were provided by the Swedish National Infrastructure for Computing (SNIC). CU was supported by grants from FWO-Vlaanderen (G0C9717N, G0C1319N) and KU Leuven (C3/19/023, C14/17/093).

